# An optogenetic approach to control and monitor inflammasome activation

**DOI:** 10.1101/2023.07.25.550490

**Authors:** Julien Nadjar, Sylvain Monnier, Estelle Bastien, Anne-Laure Huber, Christiane Oddou, Léa Bardoulet, Gabriel Ichim, Christophe Vanbelle, Bénédicte Py, Olivier Destaing, Virginie Petrilli

## Abstract

Inflammasomes are multiprotein platforms which control caspase-1 activation, leading to the processing of proinflammatory cytokines into mature and active cytokines IL-1β and IL-18, and to pyroptosis through the cleavage of gasdermin-D (GSDMD). Inflammasomes assemble upon activation of specific cytosolic pattern recognition receptors (PRRs) by damage-associated molecular patterns (DAMPs) or pathogen-associated molecular patterns (PAMPs). They converge to the nucleation of apoptosis-associated speck-like containing a caspase activation and recruitment domain (ASC) to form hetero-oligomers with caspase-1. Studying inflammasome encoding activities remains challenging because PAMPs and DAMPs are sensed by a large diversity of cytosolic and membranous PRRs. To bypass the different signals required to activate the inflammasome, we designed an optogenetic approach to temporally and quantitatively manipulate ASC assembly (*i.e.* in a PAMP- or DAMP-independent manner). We reveal that controlling light-sensitive oligomerization of ASC is sufficient to recapitulate the classical features of inflammasomes within minutes, and enabled us to decipher the complexity of volume regulation and pore opening during pyroptosis. Overall, this approach offers interesting perspective to decipher PRR signaling pathways in the field of innate immunity.

## Introduction

Inflammasomes are dynamic multiprotein complexes involved in innate immunity and inflammation. They form upon sensing of pathogen-associated molecular patterns (PAMPs) (*e.g.,* the bacterial toxin nigericin) or damage-associated molecular patterns (DAMPs) (*e.g.*, extracellular ATP) by a pattern recognition receptor (PRR), such as nod-like receptor pyrin containing 3 (NLRP3). The PRR then drives the formation of nanoclusters of adaptor-associated speck-like (ASC) necessary to activate the cysteine protease caspase-1 at the complex, through their caspase recruitment domain (CARD) (*1*). Hetero-oligomerization of the inflammasome core proteins ASC and caspase-1 filaments results in large membraneless structures, between 0.2-1 µm in diameter, called specks (Figure 1A upper panel) (*2–4*). Activated caspase-1 then cleaves proinflammatory IL-1β and IL-18 cytokines and the gasdermin-D (GSDMD) protein to induce an inflammatory form of cell death called pyroptosis (*5–7*). Indeed, the resulting GSDMD N-terminus fragments self-oligomerize and are inserted into the plasma membrane to form pores contributing to IL-1β and IL-18 release, cell swelling, membrane rupture and as a consequence lytic cell death (*8*, *9*). Different inflammasomes have been reported and are essential actors of immunity. For instance, the NLRP3 inflammasome is the most studied inflammasome and its perturbation leads to autoinflammatory disorders such as cryopyrin-associated periodic syndrome (CAPS), or metabolic disorders including gouty arthritis, atherosclerosis and type 2 diabetes (*10–14*).

**Figure 1:**
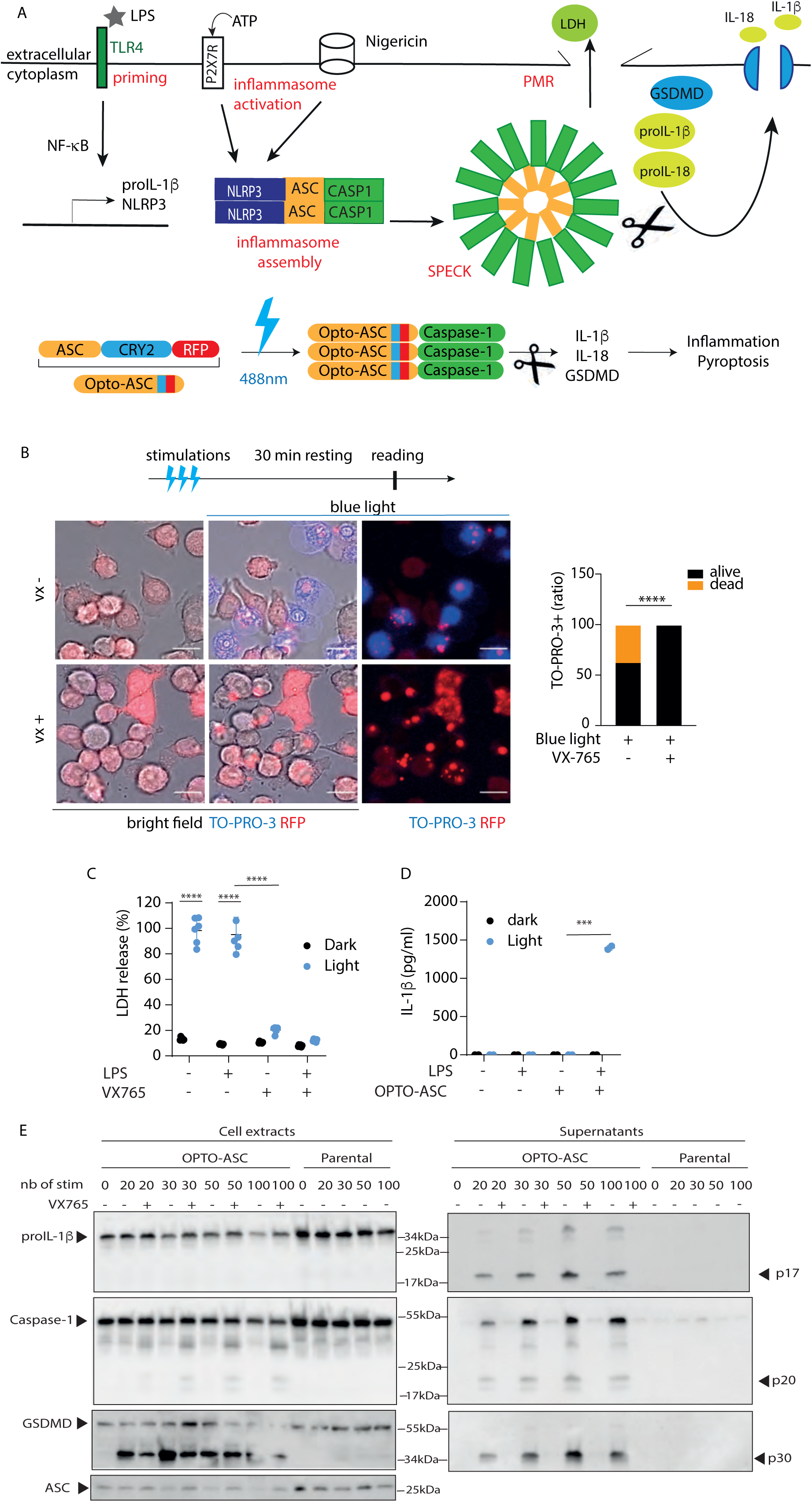
Blue light activation of opto-ASC expressing iBMDM induces caspase-1 activation, IL-1β maturation and pyroptosis. (A) Schematic representation of NLRP3 inflammasome activation followed by a diagram of the opto-ASC vector: the core inflammasome protein ASC was coupled to the blue light sensitive photolyase domain of Cryptochrome 2 (CRY2) and a RFP tag. (B) Opto-ASC iBMDM received 3 stimulations of 10 s, speck formation (RFP^+^) and cell death (TO-PRO-3^+^) were monitored 30 min post-stimulation using confocal microscopy in the absence or presence of the caspase-1 inhibitor, VX-765 (VX). Positive TO-PRO-3 nuclei were quantified in different fields. n is the number of cells, 179 < n from 3 pooled experiments. Contingency table was used for statistical analysis. Scale bar = 10 µm. (C,D) iBMDM were pre-incubated or not with LPS 0.5 ng/mL, then 50 stimulations of blue light was applied using LEDs and LDH release (C), or IL-1β secretion (D) were assessed. An unpaired t-test was used for statistical analysis, **** P<0.0001, *** P<0.001. (E) Immunoblot analysis of caspase-1 maturation, IL-1β and GSDMD cleavage upon blue light activation (ranging from 20 to 100 stimulations) of opto-ASC-expressing or parental cells (iBMDM) pre-treated with LPS in the absence or presence of VX-765 (added 30 min prior treatment). Data are representative of at least two independent experiments.

Inflammasome assembly requires stimulation with PAMPs or DAMPs, which are not exclusively engaged in inflammasome assembly, but also activate other PRRs and their downstream signaling pathways. In fact, NLRP3 inflammasome assembly also relies on an upstream priming signal, necessary to induce the expression of proIL-1β and NLRP3, and for NLRP3 post-translational modifications (Figure 1A upper panel) (*15*). This priming signal originate either from inflammatory cytokines or PAMPs, such as binding of lipopolysaccharide (LPS) to the toll-like receptor 4 (TLR4). Therefore, studying the sole output of inflammasome activation and its dynamics in cells remains challenging using classical pharmacological and genetic approaches, since signaling downstream of DAMPs or PAMPs is highly intricated and not limited to inflammasome formation. This is especially true when studying the processes of cell swelling downstream of pyroptosis activation.

To address this problem, we developed an optogenetic tool to directly control the oligomerization of the inflammasome core adaptor ASC using light, and independently of the extracellular environment. Optogenetics has been widely used in the field of neurosciences to activate neurons at the whole cell level, and is continually adapted to other cellular functions dependent on the nanoclustering of molecular complexes at the sub-cellular and cellular levels in a controlled spatial and temporal manner (*16*, *17*). Here, to spatio-temporally manipulate inflammasome activation, we engineered a new photo-oligomerizable ASC based on the CRY2 optogenetic module (*16*). We demonstrate that simple nanoclustering of ASC upon exposure to light is sufficient to induce a complete physiological inflammasome response, which is caspase-1-dependent but PAMP- and DAMP-independent. Therefore, it represents a powerful tool with many possible applications ranging from the molecular dissection of events leading to inflammasome output to the screening of specific inhibitors. As a proof-of-concept, we used this system to address cell volume regulation during inflammasome-induced pyroptosis. We precisely measured cell swelling, and unveiled a two step-mediated cell swelling before plasma membrane rupture.

## RESULTS

### Opto-ASC assembles in speck upon classical inflammasome activation

To study the output of inflammasome assembly without interference from DAMP/PAMP priming or activating signals, we designed a lentiviral vector to express the human protein ASC coupled to the photolyase homology domain (PHD) of cryptochrome 2 (CRY2) from *Arabidopsis thaliana* and to red fluorescent protein (RFP), which we named opto-ASC (Figure 1A lower panel). We reasoned that the CRY2 PHD homo-oligomerizer should initiate ASC oligomerization upon exposure to blue light, thus mimicking its physiological engagement by a PRR such as NLRP3 (*18*). Consequently, ASC would recruit and activate caspase-1 resulting in the induction of specific canonical inflammasome-dependent responses (Figure 1A lower panel).

Immortalized bone marrow-derived macrophages (iBMDM) were transduced to stably express opto-ASC (or the relevant control opto-RFP alone) (Suppl. Fig 1A) and FACS sorted according to RFP expression in order to normalize vector expression (*19*). First, we verified that the opto-ASC behaved like the endogenous protein and would not interfere with the physiological activation of inflammasomes. To achieve this, parental cell lines and cells expressing opto-ASC were primed with LPS to induce NLRP3 and proIL-1β expression, followed by activation with the bacterial toxin nigericin to trigger the assembly of the NLRP3 inflammasome (*20*). Analysis of NLRP3 inflammasome activation by immunoblot revealed that nigericin induced caspase-1 activation, in both parental and opto-ASC-expressing cells, as evidenced by the detection of cleaved caspase-1 (p20), GSDMD N terminus fragment (p30) and IL-1β (p17) in cell supernatants (Suppl. Figure 1B). A slight increase in the cleavage of these substrates was observed in the opto-ASC condition, likely due to higher expression of ASC (endogenous and exogenous ASC). Treatment with VX-765, a specific caspase-1 inhibitor, inhibited caspase-1 activation in both cell lines, clearly showing that ASC overexpression did not affect the sensitivity of the inflammasome reaction (Suppl. Figure 1B). Two other inhibitors were tested on opto-ASC-containing cells, z-VAD-fmk, a pan caspase inhibitor, and z-YVAD-fmk a caspase-1 inhibitor. Both inhibited IL-1β and caspase-1 cleavage upon treatment with nigericin (Suppl. Figure 1C). A hallmark of inflammasome assembly is the formation of ASC specks in cell cytosols (*2*, *3*). Typical ASC-RFP^+^ specks were visible upon nigericin treatment alongside plasma membrane (PM) swelling and rupture, which are features of pyroptosis (Suppl. Figure 1D). Thus, we concluded that opto-ASC assembles in specks like endogenous ASC, and that exogenous expression of opto-ASC neither affects inflammasome activation nor induces spontaneous inflammasome responses.

### Light activation of opto-ASC induces canonical inflammasome features

Next, we wondered whether blue light (488 nm) activation of opto-ASC would be sufficient to induce inflammasome assembly and activation in the absence of PAMP or DAMP stimulation. To test this, we first determined that light stimulation induced formation of Opto-ASC specks associated with pyroptosis using a confocal microscope. As expected, we observed that opto-ASC oligomerized into specks (RFP+), which was followed by cell swelling and loss of plasma membrane integrity, as illustrated by TO-PRO-3^+^ stained DNA (Figure 1B). The addition of VX-765 inhibited cell death, but not the formation of the RFP^+^ specks, indicating that light activated opto-ASC induces pyroptosis in a caspase-1-dependent manner (Figure 1B, movies 1 & 2). Interestingly, pyroptosis can better be visualized using Nanolive microscope, a cell tomography microscope, which enables users to specifically see changes in membrane, cytosol and nuclear organizations (*21*). The observation of iBMDM activated either with nigericin, or ATP or light activated opto-ASC showed that all treatments induced similar patterns of nuclear condensation and cell swelling associated with pyroptosis (see movies 3 & 4). Lactate dehydrogenase (LDH) release into the supernatant upon plasma membrane rupture (PMR) is another classical cell death marker used to quantify pyroptosis. For quantitative analysis, we activated a large population of iBMDM stably expressing opto-ASC in an incubator equipped with blue LEDs. As expected, applying light stimulation to cells stably expressing opto-ASC induced specific LDH release, independently of the presence of a priming signal (LPS), while VX-765 inhibited it (Figure 1C). Parental cells used as controls did not display an increase in cell death (Suppl. Figure 2A), demonstrating that the effect observed was not due to light phototoxicity. Thus, the simple and direct activation of opto-ASC clustering with photostimulation was sufficient to recapitulate canonical features of pyroptosis independently of any upstream signals.

To further ascertain that opto-ASC activation induced active caspase-1, we then quantified the amount of IL-1β released into the supernatant in response to blue light stimulation by ELISA. In contrast to pyroptosis, IL-1β secretion was detected only when opto-ASC activation was coupled with an LPS stimulation, since a priming signal was necessary to induce the synthesis of the proIL-1β (Figure 1D). Interestingly, our synthetic strategy allowed to uncouple the production of IL-1β from pyroptosis.

To demonstrate in a more direct manner that opto-ASC induced caspase-1 activation, we analyzed by immunoblot whether caspase-1 and its substrates were cleaved upon exposure to an increasing range of stimulations. Opto-ASC activation with twenty pulses of alternating dark/light sequences produced significant amounts of cleaved caspase-1, IL-1β and GSDMD (Figure 1E). Parental or opto-RFP-expressing cell lines used as controls did not produce any cleaved protein following light stimulation (Figure 1E and Suppl. Figure 2B). Interestingly, by applying a range of stimulations, we were able to achieve different levels of caspase-1 activation as illustrated by the increasing amounts of cleaved IL-1β and GSDMD detected in cell supernatants before reaching a plateau of substrate maturation at 50 stimulations (Figure 1E and Suppl. Figure 2C). Again, light-dependent caspase-1 activation was prevented by VX-765 treatment, confirming the physiological functions of opto-ASC activation (Figure 1E). Importantly, the different ranges of stimulation also resulted in progressive LDH release until it reached a plateau as for IL-1β (Suppl. Figure 2D). Thus, opto-ASC activity can be modulated by applying different patterns of light stimulation.

### The activity of opto-ASC relies on inflammasome effectors

Because GSDMD is the effector of pyroptosis, we transduced the opto-ASC construct in iBMDM *gsdmd*^-/-^ cells to control their response to inflammasome activation. Interestingly, light activation promoted ASC speck formation, albeit neither cell swelling nor TO-PRO-3 staining was detected unlike WT cells, indicating the absence of pyroptosis (Figure 2A). Consistently, no IL-1β was secreted (Suppl. Figure 2B). Because glycine was reported to act as a cytoprotectant by preventing PMR, we tested whether it would prevent opto-ASC-induced cell lysis (*5*). Opto-ASC-expressing cells were grown in culture medium supplemented or not with glycine. Interestingly, in WT opto-ASC cells, glycine treatment inhibited light-dependent LDH release, and therefore PMR, but not IL-1β which is secreted through GSDMD pores (Suppl. Figure 2B and E). Hence, light stimulation induces the formation of opto-ASC specks that lead to caspase-1 activation and the subsequent release of IL-1β and pyroptosis in a GSDMD dependent manner, recapitulating the canonical function of inflammasomes.

**Figure 2:**
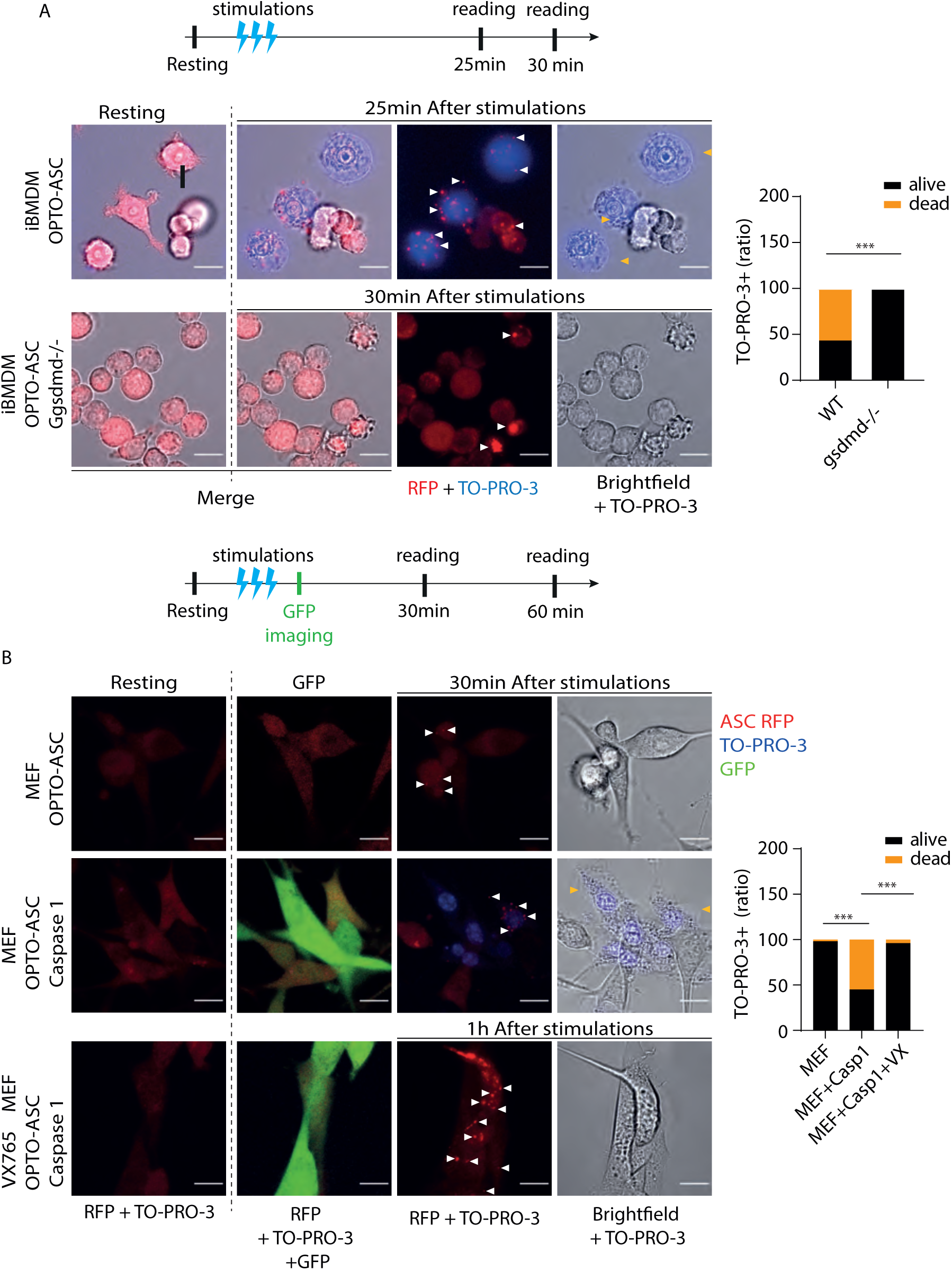
The activity of opto-ASC is GSDMD- and inflammasome-dependent. (A) WT or GSDMD-deficient iBMDM stably expressing opto-ASC were stimulated using blue light confocal; speck formation was monitored through RFP+ staining and cell death through TO-PRO-3+ nuclei staining 25-30 min after stimulation. Scale bar = 10 µm. (B) MEFs stably transduced with opto-ASC or opto-ASC + GFP-IRES-caspase-1 constructs were stimulated using blue light confocal, +/- VX-765 and speck formation was monitored through RFP+ signal and cell death through TO-PRO-3+ 30-60 min later. At least 67 cells were counted, from 2 pooled experiments. White arrows indicate ASC-RFP specks and yellow arrows indicate PM swelling. Scale bar = 10 µm. Contingency table was used for statistical analysis. *** P<0.001.

To further validate our synthetic strategy, we investigated whether we could induce inflammasome responses in mouse embryonic fibroblast (MEFs) by stably expressing opto-ASC, since they are naturally inflammasome-free cells, *i.e.* they do not express inflammasome proteins (Suppl. Figure 1A). In this cellular environment, blue light stimulation was sufficient to induce autonomous opto-ASC oligomerization, as evidenced by RFP^+^ specks, but neither cell death (absence of TO-PRO-3 staining) nor plasma membrane swelling (Figure 2B). Importantly, transducing opto-ASC alongside caspase-1 in MEFs induced pyroptosis exclusively in response to blue light stimulation, which was inhibited by the presence of VX-765 (Figure 2B) (*3*). These results clearly establish that opto-ASC does not have off-target effects and that we are able to uncouple IL-1β secretion from cell death. In addition, these results demonstrate that it is possible to use opto-ASC in an inflammasome-deficient cellular environment and to reconstitute functional inflammasome responses. This perspective is essential to control and calibrate inflammasome assays, an essential step for investigating structure-function relationships of ASC, caspase-1 or any mutants of inflammasome components.

### Controlled opto-ASC activation reveals different level of efficacy of caspase-1 inhibitors

Next, we wondered whether the opto-ASC system stably expressed in a cell type of interest could be used to compare the efficacy of pharmacological modulators of inflammasome responses in a medium throughput approach. As a proof-of-principle, we used an automated confocal microscope (Operetta CLS^TM^ microscope) to follow cell response to different settings of stimulations by modulating laser intensity or stimulation frequencies. Here, TO-PRO-3^+^ cells were used as a readout of caspase-1 activation over time (Figure 3A, D). For data quantification, we fitted the raw data (percentage of cell death) to a sigmoid function, f(x) = a/(1+exp(-k*(x-b))), and extracted the time at which 50% of the response occurred, using the inflexion point (b, the time for which half the population died) and the amplitude (a, percentage of cell death) of the response. Concerning the percentage of dead cells (a), although modulating the intensity of the laser or the number of stimulations induced cell death, the different levels of intensity only varied significantly with that at 1% and not among each other (Figure 3B). In addition, although there was an increase in the response according to the number of stimulations applied, again only the response between 1 and 2 stimulations differed significantly (Figure 3E). For the inflexion (b) point, there was no significant difference when increasing the number of stimulations or light intensity (Figure 3C and F). Thus, these results suggest that in these settings, once the energy given to the probe was sufficient to initiate the reaction, the cells underwent an all-or-nothing response (Figure 3B, C, E, F). We then applied a fixed pattern of stimulation in the absence or presence of Z-VAD-fmk, YVAD-fmk, or VX-765. We recorded TO-PRO-3+ entry as a readout of pyroptosis, and therefore of caspase-1 activation, over time. The results obtained revealed a difference in the inhibitory capacity of these different molecules with VX-765, being the most efficient at inhibiting caspase-1, whereas YVAD-FMK was the least potent at blocking cell death (Figure 3G-I). Opto-ASC is thus an efficient controllable tool for drug screening.

**Figure 3:**
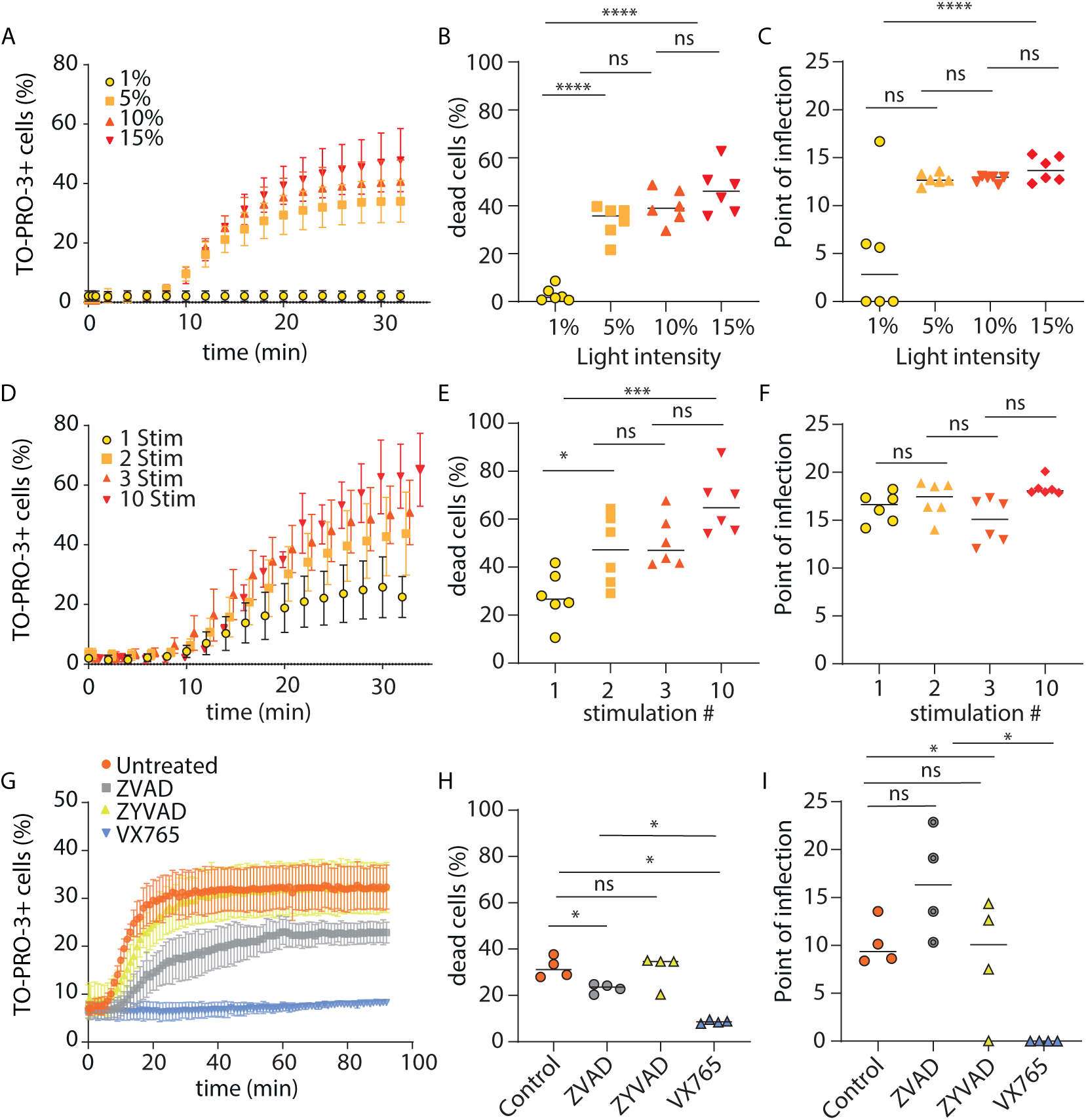
Controlled opto-ASC activation reveals different caspase-1 inhibitor efficacies. opto-ASC iBMDM were stimulated using the Operetta CLS microscope and nuclear TO-PRO-3 staining was measured over time as an indicator of cell death. (A) Different laser intensities were applied with 3 iterations or (C) different frequencies at a fixed laser intensity of 10% with 3 iterations were applied to cells. After transformation of the data from (A) using sigmoid curve, the percentage of cell death was extracted for each condition (B), as well as the inflection point (C) (as an indicator of 50% of the response). (E,F) similar extractions were performed on data from (B), the percentage of cell death was extracted for each condition (E), as well as the inflection point (F). One way anova (for a) and Mann Whitney test (for b) were used for statistical analysis. (G) The cells were stimulated using a fixed pattern, laser 10% with 3 iterations of 50 ms and cells were treated or not with different caspase inhibitors at the dose of 50 µM. (H) Percentage of cell death for conditions extracted from (G). (I) Inflection point extracted from (G). Mann Whitney tests were used for statistical analysis N=2 (N is the number of independent experiments), 4 replicates for each experiments. **** P<0.0001, *** P<0.001, * P<0.05.

### Opto-ASC reveals new aspect of cell swelling during pyroptosis

A major challenge in inflammasome research remains the precise measurement of the increase in cell volume and determining the events related to PMR during pyroptosis. One such limitation is due to the fact that many NLRP3 activators act at the level of PM by opening channels (*e.g.* ATP through the binding to P2X7R) or by permeating the PM (*e.g.* nigericin pore forming toxin) to trigger inflammasome activation (*22*). Thus, we reasoned that our optogenetic approach could be useful to precisely measure the temporal relationship between ASC activation, pore opening, cell swelling and PMR during pyroptosis by combining it with a fluorescence exclusion microscopy method (FXm) (Figure 4A) (*23*). Briefly, cells are grown in a microfluidic chamber of defined height and in the presence of cell non-permeant fluorophore-coupled dextran molecules of given sizes (Figure 4A). Images were acquired in epifluorescence with low magnification using low numerical aperture objectives (10x, NA0.32 or 20x, NA0.4). From these images, the cell volume is computed based on the exclusion of the dye (*23*). We chose to use two different sizes of dextran (10 kDa and 500 kDa) coupled to Alexa-Fluor 647 and TRITC respectively, to follow cell swelling, pore formation and PMR concomitantly. Indeed, the 500 kDa dextran (Large Dye, LD) has a hydrodynamic radius (*R_LD_* = 15.9 *nm*) larger than the GSDMD pores (*R_PORE_* = 10.75 *nm*) (*24*), whereas the 10 kDa (Small Dye, SD) has a smaller one *R_SD_* < 2 *nm* and can thus cross the pores (*25*). Pyroptosis was induced using either light activation of Opto-ASC or ATP as a standard NLRP3 inflammasome activator. Two representative examples of FXm data obtained upon pyroptosis induction following Opto-ASC activation or ATP treatment from an individual cell are shown in Figure 4B and 4C, respectively. Upon blue light activation of opto-ASC, cells started to swell until they reached a maximum volume 3.5 +/- 0.3 min later for the SD and 7.2 +/- 0.5 min for the LD (Figure 4B and Suppl Figure 3A). Then, the observed cell volume started to decrease as cells became permeable to the dyes, indicating a loss of plasma membrane integrity. Similarly, after the addition of ATP, cell swelling started and stopped 9.5 +/- 0.4 min later for the SD and 15.2 +/- 1.3 min for the LD; then as for opto-ASC activation, cells became permeable to the dyes (Figure 4C and Suppl. Figure 3B). Thus, for both conditions, the LD allowed us to determine the maximum volume reached by cells before PMR occurred, as evidenced by the peaks of the curves that were higher than the ones measured with the SD for both treatments. Interestingly, though live cell monitoring displayed similar cell swelling in the initial phase of the response for both SD and LD, the amplitude of the peaks measured with the SD were smaller since it dropped sooner than the one measured using the LD (Figure 4B,C, and Suppl. Figure 3A and B). This result indicated that the small dye entered into the cell while the process of swelling did not reach its maximum, suggesting an entry upon GSDMD pore formation.

**Figure 4:**
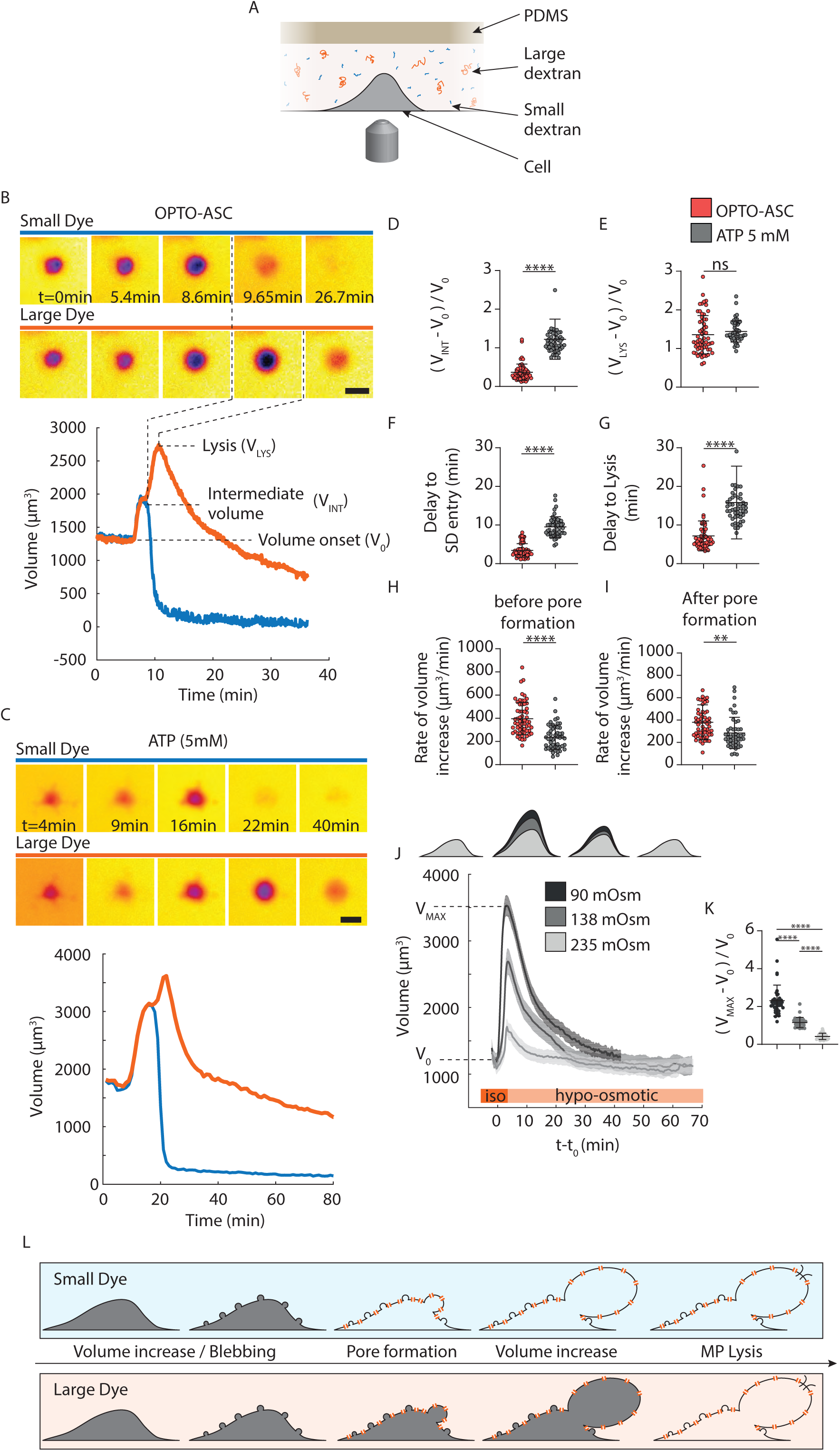
Volume increase, GSDMD pore formation and lysis are distinct events. (A) Schematic diagram of the fluorescence exclusion method (FXm) with dye-coupled dextrans of different sizes. (B) Representative volume measurement of light-induced activation of opto-ASC with large dyes (LD, red) (TRITC - Dextran 500 kDa) and small dyes (SD, blue) (Alexa647 - Dextran 10 kDa) with the corresponding time-lapse images obtained by FXm with the two different dyes. (C) Representative volume measurement of ATP (5 mM)-induced activation of WT-iBMDM cells with large dyes (LD, red) (TRITC - Dextran 500 kDa) and small dyes (SD, blue) (Alexa647 - Dextran 10 kDa) with the corresponding time-lapse images obtained by FXm with the two different dyes. Scale bars = 20 µm. (D) Mean of the intermediate volume increase normalized by the volume before increase, obtained with the SD upon light activation of opto-ASC and ATP stimulated cells. (E) Mean of the maximum volume increase normalized by the volume before increase, obtained with the LD. (F) Delay between the onset of volume increase and small dye entry for opto-ASC- or ATP-stimulated cells. (G) Delay between onset of volume increase and large dye entry. (H) Rate of volume increase for the first swelling phase (before the intermediate plateau is reached, (*V_INT_*_−_*V*_0_)/ Δ *t*). (I) Rate of volume increase for the second swelling phase before LD entry ((*V_LYS_*_−_*V_INT_*)/ Δ *t*). For D to I, N=2 and were pooled, n=59 for opto-ASC and N=2, n=50 for ATP. (J) Top: schematic diagram illustrating the increase in cell volume and adaptation measured according to the osmotic stress applied 90 (black), 138 (grey) or 235 (light grey) mOSm. Lower: Increase in iBMDM cell volume after hypo-osmotic shock (1 experiment representative of 2, n = 39, 31 and 28 cells for 90, 138 and 235 mOsm, respectively). (K) Relative maximum increase in cell volume before regulatory volume response occurred (Δ *V_max_* /*V*_0_) from data in (J). One-way ANOVA test: ** P<0.005, **** P<0.0001, mean ± S.D.. (L) Illustration of the observations described. The SD enters cells when GSDMD pores are formed while cells are still swelling, as observed with the LD. Then PMR occurs resulting in cell lysis and entry of LD.

We then compared the global ATP response to opto-ASC activation in terms of cell swelling and PMR (Figure 4D-I). Concerning the increase in relative cell volume observed with the SD, it was significantly lower upon opto-ASC activation (0.36 +/- 0.03) than upon ATP activation (1.22 +/- 0.07) (Figure 4D). Similarly, the delay to SD entry into the cells, was shorter upon opto-ASC activation (3.5 +/- 0.3 min) than upon ATP (9.5 +/- 0.4 min) (Figure 4F). However, the rate of volume increase was much faster upon opto-ASC activation in the two swelling phases (Phase 1: 400 +/- 20 µm^3^/min versus 236 +/- 15µm^3^/min for ATP; phase 2: 380+/-20 µm^3^/min versus 280 +/- 20µm^3^/min for ATP) (Figure 4H and I). The measurement using the LD revealed that, for both treatments, cells displayed a similar swelling before lysis (Δ*V* /*V*_0_ = 1.35 +/- 0.1 for opto-ASC activation and 1.45 +/- 0.1 for ATP) (Figure 4E), whereas the delay to cell lysis and the rate of volume increase was faster upon opto-ASC stimulation (Figure 4G and I). These results show that ATP by itself also affects the cell volume upstream of inflammasome activation and are in accordance with the fact that ATP binding to P2X7R opens a cation-selective channel (*26*). In addition, these results highlight that upon GSDMD pore formation, cells kept swelling to double in size before PMR occurred. Hence, our opto-ASC construct enabled us to uncouple the signaling events upstream of inflammasome activation (P2X7R opening) from the downstream effects of caspase-1 activation to measure events exclusively related to cell swelling, GSDMD pore formation and PMR.

Our result showed that during pyroptosis cells double in size. PMR downstream of inflammasome activation has often been suggested to originate from PM mechanical stress following an osmotic imbalance due to pore formation (*27*). We therefore compared the response of iBMDM to an osmotic shock with the one induced upon pyroptosis. To that extent, three hypo-osmotic stresses ranging from 90 to 235 mOsm were applied to cells to induce passive swelling, and volume adaptation was recorded (Figure 4J, K). The data obtained showed that the 3 stresses induced moderate to large increases in cell volumes, but not PMR, as cells returned to their original size after swelling through regulatory mechanisms (*28*). The maximum volumes reached were in accordance with theoretical predictions from the Ponder – Van’t Hoff model and gives an osmotically inactive volume for iBMDM cells of 18.6% in agreement with our expectations (See Suppl. Figure 3C) (*29*, *30*). More importantly, the 90 mOsm stress tripled the size of cells without inducing PMR, demonstrating that iBMDM have a sufficient amount of quickly available PM to sustain large deformations without undergoing PMR (Figure 4K). Since inflammasome activation using opto-ASC, induced PMR when cells reached twice their volume, we concluded that the PMR associated with pyroptosis is not a passive process due to osmotic shock but is most likely a regulated process that promotes plasma membrane weakening.

## Discussion

Inflammasome formation results in three main biological outcomes: the formation of speck, the maturation and secretion of inflammatory cytokines, and pyroptosis. Here, we generated a new specific photoactivable inflammasome, opto-ASC, based on optogenetic methods using the photosensitive element CRY2 which was designed to activate the inflammasome in a PAMP- and DAMP-independent manner. We provide proof–of-concept that an optogenetic approach is a powerful tool suitable to control and dissect inflammasome signaling thanks to its fast and reversible activation. This approach is therefore more powerful than other reverse engineering systems based on chemical dimerization (*31*). We demonstrated that we were able to uncouple cytokine production from pyroptosis and to reconstitute inflammasome-deficient cells with minimal components, thus this tool is relevant to many cell types of interest. In addition, the opto-ASC system could also be useful to explore encoded signals involved in inflammasome activation using genetic or chemical screens. While preparing this manuscript, the team of P. Broz reported a similar optogenetic approach to directly control caspase activation, and also demonstrated that it was suitable to induce pyroptosis (*32*).

An optogenetic approach does not only offer fast activation but also reversible activation. Indeed, 4-6 min post-illumination, the CRY2 complex dissociates, and this property is frequently used to study the dynamics of complex dissociation (*33*). Hence, 4-6 min after stimulation, ASC specks could be predicted to dissociate. However, here, in conditions in which pyroptosis was prevented by adjunction of a caspase-1 inhibitor or by working in inflammasome-deficient cells, we observed that ASC specks did not dissociate after more than 10 min post-stimulation, supporting the notion that ASC behaves like a prion protein *in vivo* due to intrinsic oligomerization properties of its PYD and CARD (*18*). Thus, opto-ASC could be useful to study speck assembly, binding properties of specific ASC mutants or to discover novel functions unrelated to the inflammasome.

To our knowledge, our study is the first to report with such precision the cell volume increase during pyroptosis. We unraveled that the cell swelling caused by the sole caspase-1 activation is about 100% of the initial cell volume, whereas following classical osmotic stress, the cell volume increased by about 300% without undergoing any PMR. Thus, during pyroptosis, cell swelling and PMR are tightly coupled and regulated mechanisms. On the contrary to the proposed model by Davis et al, suggesting that *in vitro* pyroptotic cells remain swollen, and that PMR occurs via mechanical stress like shear stress, we demonstrated using a large dextran that inflammasome activation induces PMR after the swelling phase (*27*). Our results are consistent with recent findings from Dixit and colleagues who described that PMR is an active mechanism driven by Ninjurin-1 activation (*34*). The opto-ASC tool also revealed that during pyroptosis, cell swelling occurred in two phases, with an intermediate plateau that most likely reflects the formation of the GSDMD pores (Figure 4L). Importantly, future quantitative analyses using opto-ASC should unveil the molecular mechanisms controlling the GSDMD pore formation and the biophysics of PMR. Finally, we hope that this type of approach will open new perspective in the field of innate immunity to refine PRR signaling pathways.

## Material and method

### Reagents

z-YVAD-fmk and z-VAD-fmk were purchased from Bachem. TO-PRO™3 Ready Flow™ Reagent and Alexa-Fluor 647-Dextran 10,000 Da was from Thermofisher and TRITC-Dextran 500,000 Da from Tdb Labs (Sweden) and Polydimethylsiloxane (PDMS) Sylgard 184 from Dow Corning. Nigericin (N7143) was from Sigma Aldrich. Ultra-pure LPS (*Escherichia coli* 0111:B4), ATP (tlrp-ATP) and VX-765 were purchased from Invivogen.

### Cloning of opto-ASC construct and stable expression in immortalized macrophages

In this study, human ASC, CRY2 and TagRFP were cloned into a pSico R vector (Addgene 11579) digested with Nhe1/Eco R1. The CAG promoter was then cloned into this vector in order to moderate overexpression of opto-ASC and make its expression less sensitive to methylation processes occurring in numerous primary cell types. The pSico-CAGpromoter-CRY2-TagRFP backbone (sequence available in the Supplementary table) was thus obtained by amplification with PHUSION high fidelity DNA polymerase (NEB) and using Gibson assembly (NEB) following the supplier’s instruction and the indicated primers: primer asc fwd = ttagtgaaccgtcagatccgctagcATGGG GCG CGC GCG CGA C, primer asc rvs = ccatcttcatcttaa t taa GCTCCGCTCCAGGTCCTCCACC, primer cry 2 fwd = ggagcggagcttaattaaGATGAAGATGGACAAAAAGACTATAG, primer cry2 rvs = cttaat c a g c t c g c t c a t t t c g a a T G C T G C T C C G AT C AT G AT C, p r i m e r Ta g R F P f w d = GATCATGATCGGAGCAGCATTCGAAATGAGCGAGCTGATTAAG, primer Tag RFP rvs = agttattaggtccctcgacgaattctcaCTTGTGCCCCAGTTTGC, primer CAG fwd = agtactaggatccattaggcggccgcGAGTTCCGCGTTACATAACTTAC, primer CAG rvs = cgtcgcgcgcgcgccccatgctagTGATGAGACAGCACAATAAC

### Lentivirus production, cell infection and sorting

Lentiviruses were produced by co-transfecting pC57GPBEB GagPol MLV, pSUSVSVG and each plasmid of interest using lipofectamine 2000 (Invitrogen) in HEK293 FT (precious gift of Dr Nègre from the ANIRA platform) cells plated in 6-well plates at 50% confluency. Medium was changed 24 h later. The viral supernatant was collected 72 h later and filtered using 0.45 μm filters. Macrophages were plated in 6-well plates at 60% confluency the day of infection. The filtered supernatant was directly used to infect cells of interest. The medium was changed 24 h after infection. After 10 days, cells were FACS sorted (Aria cell sorter 2000, or Aria III using Diva software (BD Biosciences) based on the level of expression of RFP-tagged opto-ASC using 561 nm laser.

### Retroviral transduction

The plasmid encoding GFP-IRES-Caspase-1 was obtained from P. Broz (*3*, *35*). To produce retroviral particles, Phoenix-Eco packaging cells were co transfected with pMSC2.2-expressing vectors for caspase-1 wild type using lipofectamine 2000 reagent (life technologies). The medium was removed after 24 h and replaced with fresh complete medium. After 48 h the medium was filtered twice using 0.45µm filters. MEFs were transduced, isolated and cloned on FACS Aria III (BD Biosciences) using Diva software (BD Biosciences).

### Cell lines

Murine iBMDM (immortalized bone marrow-derived macrophages), MEFs (mouse embryonic fibroblasts), HEK-293T (human embryonic kidney 293T cells), PlatE (Platinum E cell line) were cultured with DMEM 4.5 g/L glucose (Gibco) supplemented with 10% FBS, 1% penicillin-streptomycin, 1% Glutamax, 1% sodium pyruvate (all Invitrogen). Caspase inhibitors (50 µM) were added 30 min prior to stimulation.

### Live imaging of pyroptosis process

Cells were seeded into Lab-Tek II chamber Slide (Nunc, Roskilde, Denmark) in phenol-red free medium, 24 h before imaging. Cellular viability was assessed using TO-PRO-3 Ready Flow™ Reagent (dilution: 1/2000). Optogenetic model activation and pyroptosis process imaging was performed in controlled atmosphere 37°C, 5 % CO2, with confocal fluorescence microscope LSM-880 Zeiss (Objective 40x NA 1.3). Cells were illuminated 3 times every 10 seconds with 488nm laser (Light dose 45 mJ/cm²). 30 min later, speck formation (RFP) and dying cells were imaged using 561 and 633 nm lasers, respectively. Image analysis was performed with Fiji software. For live-cell tomography images acquisition, cells were seeded on ibidi plates 24 h before imaging with Nanolive 3D Cell Explorer-Fluo.

### IL-1**β** and LDH release analysis

48h after plating, cells were primed or not with LPS 0.5 ng/ml for 3 h and treated or not with caspase-1 inhibitor (VX-765) 50 μM for 30 min in optiMEM serum-free medium. Cells were illuminated or not 50 times with 0.8s flash using an incubator equipped with blue LED (1.8 mW) as initially described by the lab of Janovjak (*36*) (Light dose 42 mJ/cm²). 1 h after illumination, dosage of IL-1β and LDH released were quantified on cell supernatants with a mouse IL-1β/IL-1F2 DuoSet ELISA (R&D Systems) and CytoTox 96 Non-Radioactive Cytotoxicity Assay (Promega, France), respectively.

### Gradual activation of opto-ASC cells with the Operetta CLS

Death induction followed by light illumination was monitored at 37°C, 5% CO2 using TO-PRO-3 for cell viability and a Perkin Elmer Operetta CLS High-Content Analysis System (20x water-objective) controlled by Harmony software (Perkin Elmer, Germany). Light activation was performed with 488 nm LED of Operetta microscope. Opto-ASC expressing iBMDMs were activated by 3 light stimulations of increasing LED intensity (1, 5, 10 and 15 % of LED power) or an increasing number of light stimulations with a fixed intensity of 10 %. For the assessment of caspase-1 inhibitors activities, cells received 3 stimulations of 10% intensity. Image quantification was conducted with Columbus software (Perkin Elmer, Germany).

#### Power density table

| LED power (%) | fluency (mJ/cm<sup>2</sup>) | Nb of stim | fluency (mJ/cm<sup>2</sup>) |
| --- | --- | --- | --- |
| 1 | 2,851675579 | 1 | 75,25254999 |
| 2 | 16,63477421 | 2 | 150,5051 |
| 5 | 102,1850416 | 3 | 225,75765 |
| 10 | 225,75765 | 10 | 752,5254999 |

### Fabrication of FXm chips

Chips were made by pouring PDMS elastomer and curing agent (1:10, Sylgard 184 from Dow Corning) on a mold and was cured for at least 2 h at 65°C. Molds were made on a silicon wafer with SU-8 photoresist using classical photolithography techniques. 3 mm inlets and outlets in the PDMS were punched and the PDMS chips were cut to fit on glass coverslips. The coverslips were bond onto 35 petri dishes with Norland Optical Adhesive 81 (Norland) for 1 minute at UV 324 nm. Chips were bonded to glass coverslips by exposure to oxygen plasma for 30 s, and dried for 5 min at 65°C immediately after bonding. These systems could then be stored for a week at +4°C and washed with PBS prior to seeding cells.

### Volume measurement with FXm

Cell volumes were obtained using the fluorescence exclusion method (FXm) as detailed in (Zlotek-Zlotkiewicz *et al*, 2015). Briefly, 5×10^4 cells were plated on PDMS chip with medium a day before experiments. ATP-treated cells were primed with LPS. For volume measurement, phenol red free medium was supplemented with a fluorescent dye coupled dextran molecules. Fluorescence was thus excluded by the cells and cell volumes were obtained by integrating the fluorescence intensity over the cell. Excitation and acquisition were performed at 37°C in CO_2_-independent medium (Life Technologies) supplemented with 1 g/L Alexa Fluor 647 dextran (10 kDa) and TRITC dextran (500 kDa) using an epifluorescence microscope (Leica DMi8) with a 10x objective (NA. 0.3, LEICA). Light activation was performed with 488 nm LED (25% of a PE300 LED from CoolLED) during 10 s (Light dose 3720 mJ/cm².

For osmotic shocks experiments, hypotonic solutions were generated by addition of dH20 to imaging medium. Then, Alexa Fluor 647 dextran (10 kDa) resuspended in PBS was added at a final concentration of 1 g/L.

Analyses were performed using homemade Matlab scripts (MathWorks).

### Analysis of inflammasome activation by Western blot

As described by Guey et al, with the following modification, semi-dry transfer was carried using the Transblot turbo system from Biorad (*37*).

#### Antibodies

|  |  |  |
| --- | --- | --- |
| ASC | Ab AL177 1:1,000 | Adipogen |
| tRFP | AB233 1:1,000 | Evrogen |
| Caspase-1 | A G-20 B-0042 (Casper),<br>1:1,000 | Adipogen |
| IL-1β | AF-401-NA 1:1,000 | R&Dsystems |
| GSDMD | EPR19828 ab209845, 1:1,000 | Abcam |
| actin | C4, 1:50,000 | ICN-MP |
| tubulin | CP06 1:10,000 | Millipore |
| Rabbit | Peroxidase anti-rabbit<br>715-035-150 | Jackson ImmunoResearch |
| Mouse | Peroxidase anti-mouse<br>711-035-152 | Jackson ImmunoResearch |
| Goat | Peroxidase anti-goat<br>705-035-147 | Jackson ImmunoResearch |

### Statistical analysis

Data are expressed as mean ± SD. The statistical significance between experimental conditions was determined by contingency table, unpaired t-test with two tail or Mann Whitney test. All data were analyzed with GraphPad Prism 9.0 (San Diego, CA, USA). **** P<0.0001, *** P<0.001, * P<0.05.

## Acknowledgements

Funding: Ligue Nationale contre le cancer comité de l’Ain (VP), La Ligue Nationale Contre le Cancer (GI), LLNC as “Equipe labellisée Ligue 2014” (EL2014.LNCC) (OD), Fondation de France (GI), fondation FINOVI (BP, VP), ANR-22-CE15-0032-01 (VP and BP), Institut Convergence PLAsCAN, ANR-17-CONV-0002 (VP, SM), Fondation pour la Recherche Médicale as « équipe labellisée » DEQ20170336744 (VP) and DEQ20170336702 (OD), LabEx DEVweCAN (University of Lyon, GI), CLARA cancéropole (VP), Agence Nationale de la Recherche (ANR) Young Researchers Project ANR-18-CE13-0005-01 (GI), ANR-13-JSV2-0003-01(OD), ANR-19-CE13-0030 (EB), ERC-2013-CoG_616986 (BP), Marie Sklodowska-Curie fellowship n°751216 (ALH), the fondation Line Pomaret Delalande PLP202110014593 (LB).

We thank Pr ES Alnemri for sharing iBMDM cells, B. Manship for English editing of the manuscript and A. Tissier for critical reading of the manuscript.

The authors declare no conflict of interest.

---

**Supplementary Figure 1:**
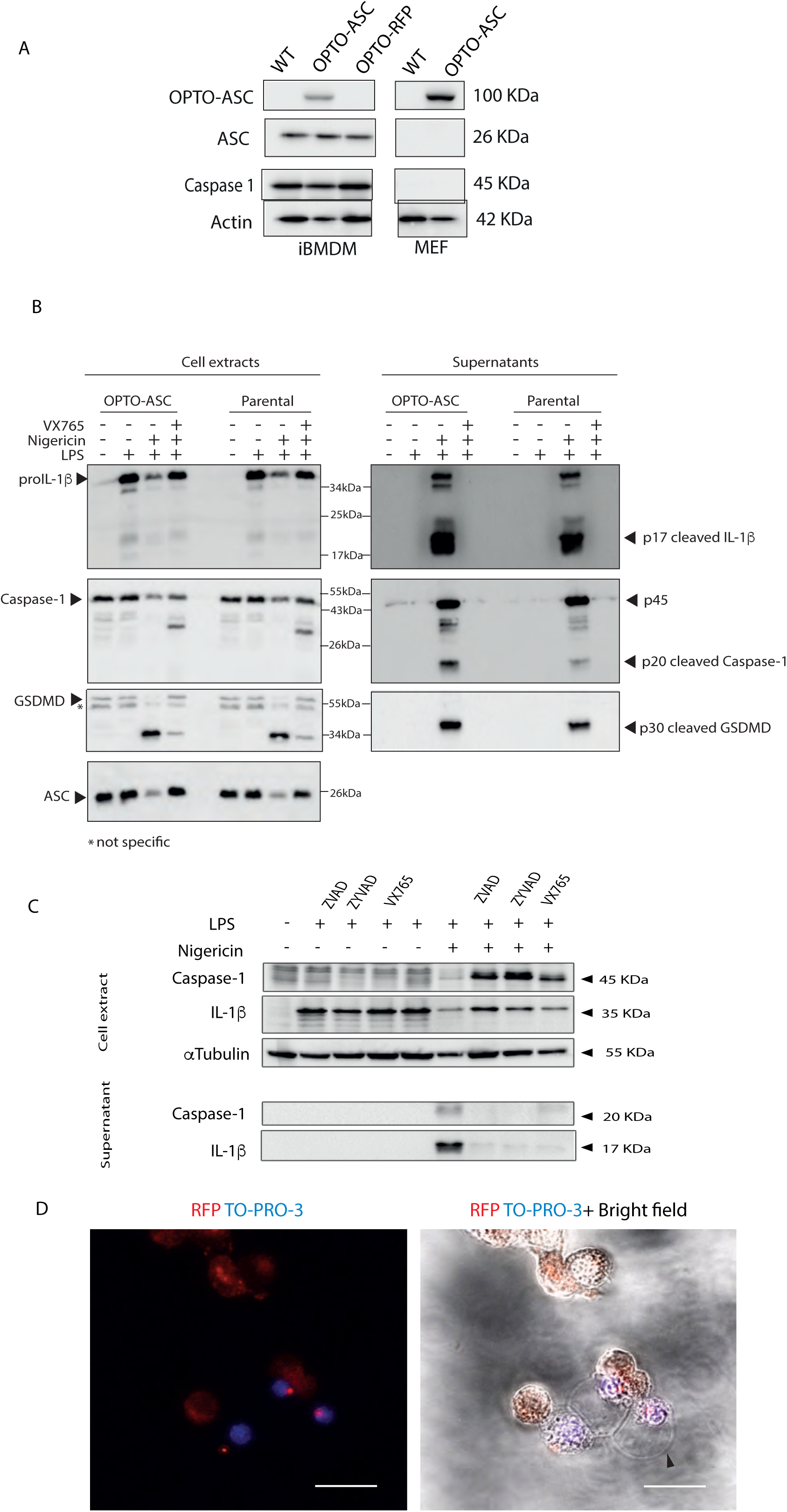
Validation of the opto-ASC vector. (A) Immunoblot showing the level of opto-ASC and inflammasome protein expression in iBMDM and MEFs. Actin is used as a loading control. (B) Immunoblot analysis of NLRP3 inflammasome activation in iBMDM cells expressing or not opto-ASC. Cells were primed with LPS 0.5 ng/mL for 3 h followed by nigericin (Nig) treatment 10 µM 3 h, in the presence or absence of the caspase-1 inhibitor VX-765 (30 min prior Nig). (C) Immunoblot showing the effect of Z-VAD, Z-YVAD, and VX-765 on caspase-1 activation upon exposure to LPS+Nig in iBMDM-expressing opto-ASC. Representative of at least 2 experiments. (D) Confocal images illustrating the formation of ASC-RFP^+^ speck and pyroptosis bleb (black arrow) after LPS+Nig treatment in opto-ASC iBMDM cells x40. Representative of at least 2 independent experiments.

**Supplementary Figure 2:**
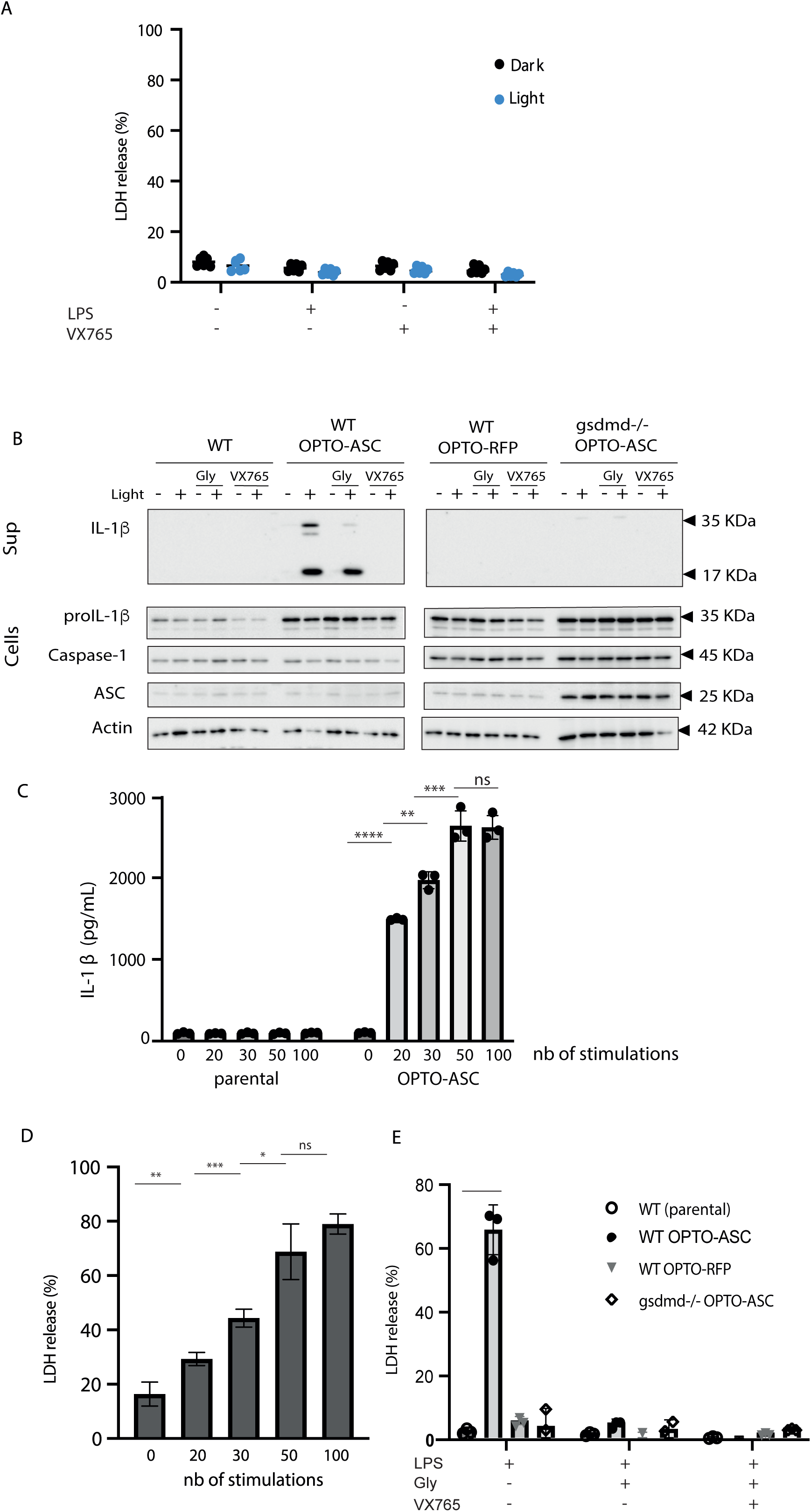
opto-ASC activation induces IL-1β release in a GSDMD-dependent manner. (A) LDH release by parental cells (iBMDM) stimulated as described in Figure 1C. (B) Parental iBMDM (WT), Opto-ASC-expressing WT iBMDM, OPTO-RFP-expressing WT iBMDM and *gsdmd*-/- iBMDM expressing opto-ASC were pre-treated with LPS then activated or not using 50 mW/cm^2^ LED stimulation in the presence or absence of glycine (20 mM) or in the presence or absence of VX-765 (50 µM). IL-1β, caspase-1 and GSDMD cleavage and release into the cell supernatant were analyzed by immunoblot (1 out 2 representative experiment). (C) LDH assay performed in the same conditions as in Figure 1E (1 out 2 representative experiment). Unpaired t test was used for statistical analysis (D) IL-1β measurement in cell supernatant from Figure 1E. One representative experiment out of 2. **** P<0.0001, *** P<0.001, ** P<0.01, * P<0.05.

**Supplementary Figure 3:**
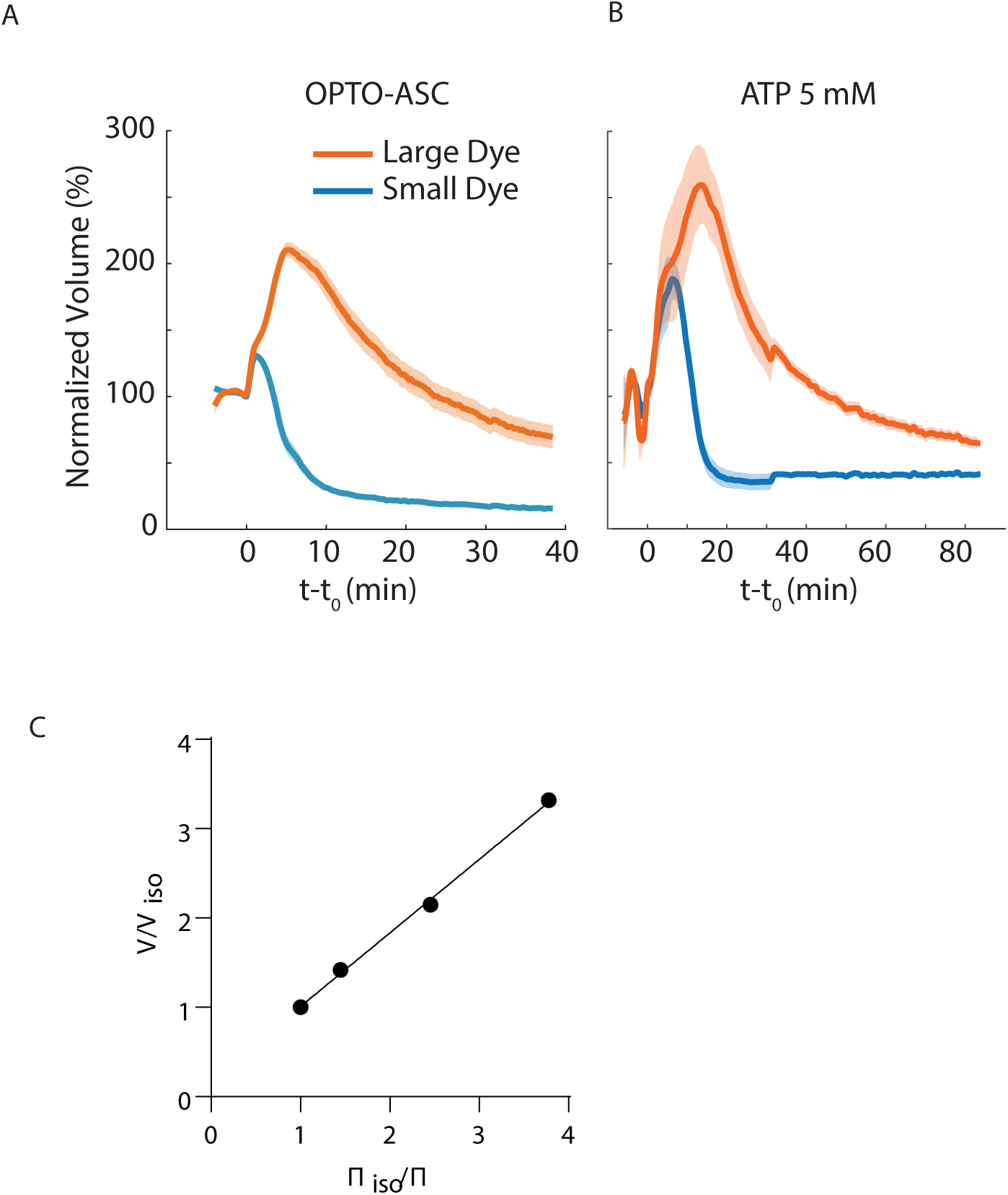
Mean volume trajectories and Ponder – Van’t Hoff plot. (A,B) Mean volume trajectories for opto-ASC (A) and ATP (5 mM) (B) upon induction of pyroptosis corresponding to panels in Figure 4B and C. N=2, n=59 for opto-ASC and N=2, n=50 for ATP. (C) Plot of maximum volume reached after hypo-osmotic shock and fitting of the data using the Ponder – Van’t Hoff equation 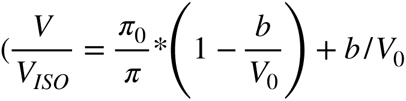 with b being osmotically-inactive volume (*b* = 18.6*%*).

**Movies 1 and 2:** confocal images over time of iBMDM cells expressing opto-ASC that were primed with LPS 3h then treated with nigericin 3h in absence (video 1) or in presence of VX765 (video 2). Scale bar = 10 µm

**Movies 3 and 4:** Nanolive images over time of iBMDM cells expressing opto-ASC that were stimulated with light or primed with LPS and treated with nigericin (video 3), or stimulated with light or primed with LPS and treated with ATP (video 4). Scale bar = 10 µm

**Supplementary table:**
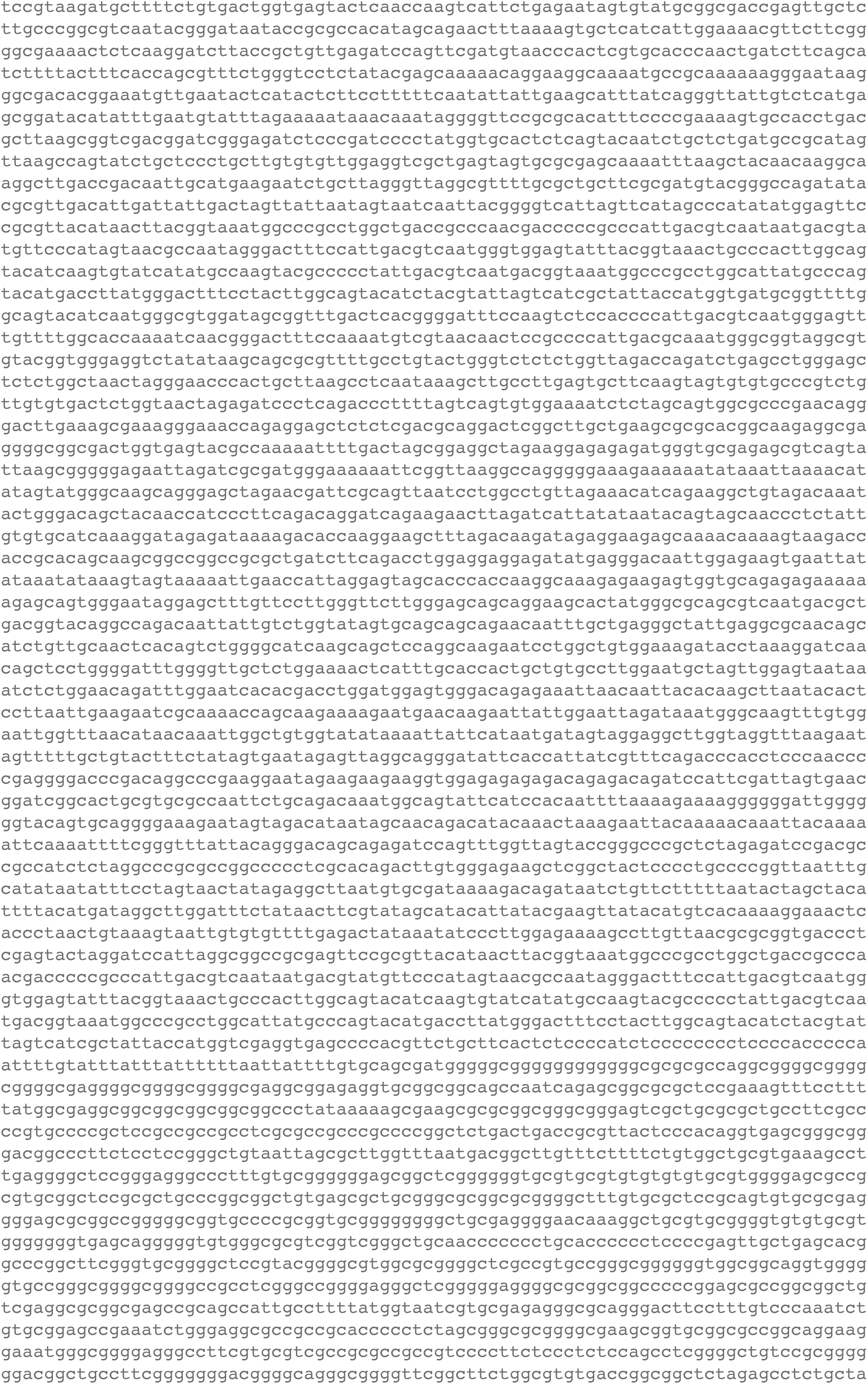
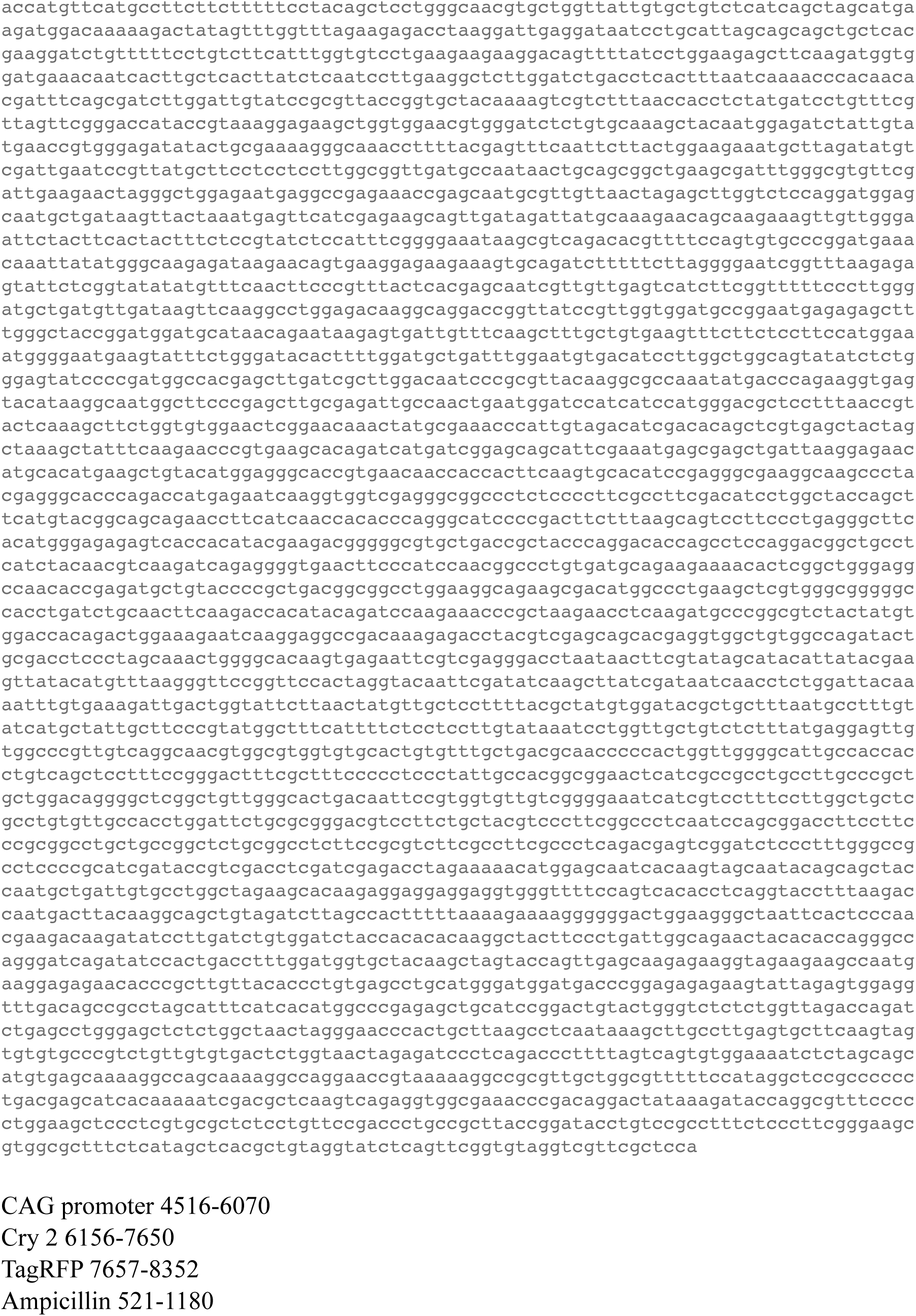
vector backbone.

## Notes

### Competing Interest Statement

The authors have declared no competing interest.

